# Genetic Sex Validation for Sample Tracking in Clinical Testing

**DOI:** 10.1101/2021.12.13.470986

**Authors:** Jianhong Hu, Viktoriya Korchina, Hana Zouk, Maegan V. Harden, David Murdock, Alyssa Macbeth, Steven M. Harrison, Niall Lennon, Christie Kovar, Adithya Balasubramanian, Lan Zhang, Gauthami Chandanavelli, Divya Pasham, Robb Rowley, Ken Wiley, Maureen E. Smith, Adam Gordon, Gail P. Jarvik, Patrick Sleiman, Melissa A Kelly, Sarah T. Bland, Mullai Murugan, Eric Venner, Eric Boerwinkle, the eMERGE III consortium, Cynthia Prows, Lisa Mahanta, Heidi L. Rehm, Richard A. Gibbs, Donna M. Muzny

## Abstract

**Background:** Next generation DNA sequencing (NGS) has been rapidly adopted by clinical testing laboratories for detection of germline and somatic genetic variants. The complexity of sample processing in a clinical DNA sequencing laboratory creates multiple opportunities for sample identification errors, demanding stringent quality control procedures.

**Methods:** We utilized DNA genotyping via a 96-SNP PCR panel applied at sample acquisition in comparison to the final sequence, for tracking of sample identity throughout the sequencing pipeline. The 96-SNP PCR panel’s inclusion of sex SNPs also provides a mechanism for a genotype-based comparison to recorded sex at sample collection for identification. This approach was implemented in the clinical genomic testing pathways, in the multi-center Electronic Medical Records and Genomics (eMERGE) Phase III program

**Results:** We identified 110 inconsistencies from 25,015 (0.44%) clinical samples, when comparing the 96-SNP PCR panel data to the test requisition-provided sex. The 96-SNP PCR panel genetic sex predictions were confirmed using additional SNP sites in the sequencing data or high-density hybridization-based genotyping arrays. Results identified clerical errors, samples from transgender participants and stem cell or bone marrow transplant patients and undetermined sample mix-ups.

**Conclusion:** The 96-SNP PCR panel provides a cost-effective, robust tool for tracking samples within DNA sequencing laboratories, while the ability to predict sex from genotyping data provides an additional quality control measure for all procedures, beginning with sample collections. While not sufficient to detect all sample mix-ups, the inclusion of genetic versus reported sex matching can give estimates of the rate of errors in sample collection systems.

## Background

To date, there are few published studies that systematically describe sample tracking methods and sample misidentification rates in laboratory samples. One study identified that, while all elements of clinical laboratory workflows can be targeted for quality improvement, the pre-analytical phase is the most vulnerable part of the testing procedure and should be considered to be the most common place for errors to occur. It was reported that more than 46% of all errors occur during the pre-analytical phase and were caused by inappropriate test requests, order entry errors, patient misidentification, and labelling errors, while less than 10% of the errors happened during the testing analytical processes(1).

Next generation DNA sequencing (NGS) technologies are capable of sequencing hundreds of samples simultaneously and have been widely implemented in clinical laboratories for high-throughput detection of germline and somatic genetic mutations (2–4). A clinical NGS test can be designed to sequence a panel of selected genes, or the entire exome/genome with a generally similar workflow. Next generation sequencing typically consists of three general phases: (i) the pre-analytic phase including sample collection, DNA extraction and shipment; (ii) the analytic phase of NGS library preparation, DNA sequencing, bioinformatics analysis of sequencing data, and variant verification; and (iii) a post-analytic phase including clinical report generation and delivery. At large scale, the sample processing in each of these phases is inherently subject to sample tracking and identification errors. Validation and tracking of sample identity therefore is a basic and important aspect of effective clinical NGS testing.

Natural DNA sequence variation can be used to distinguish different human samples with high accuracy and affords the opportunity to improve sample tracking in clinical laboratories. Two genotyping methods are commonly used for individual human sample identification in genetic testing: short tandem repeat (STR) profiling and single nucleotide polymorphism (SNP) analysis. Short tandem repeats are variable repeated segments of DNA that are typically 2-6 base pairs in length, scattered throughout the genome. Short tandem repeat profiling methods can employ multiplex PCR to simultaneously assay up to 16 loci(5). Although STR testing is the most common method for identification of DNA samples, it is not ideal in the context of clinical NGS testing, as the STRs are located in non-coding regions, prone to high sequencing error rates, and often require longer than typical sequencing read lengths to precisely define the number of repeats. In contrast, SNP genotyping is technically relatively simple and can be directed at many specific locations in the genome, including coding exons of genes. In contrast to STRs, SNP profiling fits naturally into the NGS platform as the SNPs can be tested by allele-specific PCR, array hybridization or DNA sequencing. The application of SNP-based sample validation and tracking based upon either polymerase chain reaction (PCR) amplification or hybridization, has recently been reported in NGS studies(6, 7).

In this study, a 96-SNP PCR panel was used to trace samples through the clinical NGS workflow in the United States National Institute of Health’s Electronic Medical Records and Genomics Phase III (eMERGE) program. The eMERGE Phase III program aimed to advance medical care by coordinating and integrating genomic characterization of patient samples with electronic medical records. It combines DNA biorepository data of predominantly healthy individuals with electronic medical record (EMR) systems for large-scale, high-throughput genetic research to determine clinically actionable findings(8). The eMERGE Phase III network linked together 11 sample collection sites and 2 clinical genetic testing laboratories: the Human Genome Sequencing Center Clinical Laboratory (HGSC-CL) at Baylor College of Medicine (BCM); and the Mass General Brigham (previously Partners HealthCare) Laboratory for Molecular Medicine (LMM) in partnership with the Clinical Research Sequencing Platform (CRSP) at the Broad Institute of MIT and Harvard. A total of 25,015 clinical DNA samples from volunteers enrolled in the eMERGE Phase III network were sequenced. The performance evaluation reported here shows that the 96-SNP PCR panel-based procedure provided a robust method for sample tracking in the clinical NGS workflow and that the testing of sex can provide a valuable quality control tool that reveals sample mislabeling or entry errors at sample collection sites, prior to NGS testing.

## Methods

### DNA SNP Test Sites and the Fluidigm SNP genotyping assay

The DNA testing for eMERGE III was carried out in two clinical laboratories that harmonized their methods for the program(8). For sample tracking, each testing laboratory utilized a 96-SNP PCR panel based upon the technology of the Fluidigm^®^ SNPtrace™ Panel Genotyping Assay (“SNPtrace™ Panel”) but incorporated different selected SNPs to enable sample identification and ancestry. Each 96-SNP PCR panel contained one subset of SNPs that were located on the sex-chromosomes and a second subset that was chosen to also be within the region of the genome targeted by the gene sequencing capture assay used in the eMERGE Phase III Network program, to analyze 20,015 participant’s DNA samples. The overall workflow, including sample accession, genotyping and sequencing, and reconciliation of the data from each source, is shown in Figure One. Details of the SNP content for the version of the Fluidigm SNP-PCR Panel assay employed at each site are shown in Supplementary material.

The 96-SNP PCR panel utilized at the BCM-HGSC-CL was essentially the Fluidigm® SNPtrace™ Panel Genotyping Assay (“SNPtrace™ Panel”), replacing 19 of the original 96 sites to match the regions of the genome that were specifically targeted in eMERGE III. The remaining SNP sites were those provided by the vendor and included sites that could be used to infer sex and ethnicity(9, 10). Overall, there were 90 autosomal SNPs and 6 sex chromosome SNPs, including 3 on Chromosome X and 3 on Chromosome Y. At the Broad Institute, the chosen SNPs included 95 autosomal SNPs and 1 sex determining assay SNP, covering the AMELX and AMELY gene (AMG_3B) which has a 6 base pair insertion for male and a 6 basepair deletion for female. The panel was designed to have overlapping SNPs with a variety of the Broad Institute’s internal sequencing and genotyping assays, which enables genotype and sex concordance checking to occur across a variety of workflows.

The 96-SNP PCR panel genotyping assays were performed according to the manufacturer’s (Fluidigm) recommendations as in the SNPtrace Panel User Guide. In brief, 12.5 ng of genomic DNA was amplified, first with a pool of 96 specific target amplification (STA) primers and 96 locus specific primers (LSP), to enrich the 96 target SNP loci. Up to 96 samples are processed in a batch on a 96-well plate. Samples were thermal-cycled according to manufacture’s protocol. After priming of 96.96 Dynamic Array™ Integrated Fluidic Circuit (IFC) in the IFC Controller HX, the STA amplified genomic DNA and the SNPtrace Assay Primer Mix which contains the Allele Specific Primers (ASPs) and the LSP were pipetted in the sample inlets and assay inlets of the IFC, respectively. Samples and assays were loaded in the IFC Controller HX and thermal cycled within the BioMark HD System with SNPtype 96×96 v1 program. The endpoint images were captured with BioMark HD System and analyzed with Fluidigm SNP Genotyping Analysis Software. A ‘per sample’ cutoff for the SNP call rate was established at 90%, for sample pass or fail.

Automated interpretation of sex was generated by the vendor software, which required ‘female’ assignments to have heterozygous/homozygous X-chromosome SNPs and no ‘Y’ SNP assignments. Conversely, ‘male’ assignments were required to have both X and Y SNP genotypes.

### Illumina Infinium SNP array assays

Illumina Infinium SNP array assays were performed with Illumina HumanCoreExome v1-3 BeadChips (Cat. No. 20024662) according to the manufacturer. The HumanCoreExome BeadChip contains 500K genomic variant sites, including more than 12,900 located on the X chromosome, that are informative for genetic sex prediction. Genotyping was by the manufacturer’s methods. Genomic DNA (200 ng) was denatured and then isothermally amplified, followed by enzymatic fragmentation and hybridization. After hybridization, washing steps were conducted to remove non-specifically bound DNA fragments and an allele-specific single-base extension reaction was utilized, to incorporate labeled nucleotides into the bead-bound primers. A staining process was performed to amplify signals from the labeled extended primers. The coated BeadChips were imaged using the Illumina iScan system. SNP calls were collected using the GenomeStudio Version 2011.1 software (Illumina Inc.). SNP clustering and genotyping were performed with default settings recommended by the manufacturer. A cutoff for the SNP call rate was 90% and data from samples passing this metric were used for analyses.

### Next Generation Sequencing Library Preparation and Capture Sequencing

DNA sequencing using a custom oligonucleotide DNA capture-enrichment panel for the eMERGE phase III program has been described previously(8). The eMERGE capture design, CAREseq, targets approximately 550 kb of the human genome, including 59 genes recommended for testing by the American College of Medical Genetics (ACMG-59)(11). The targeted region also contains 1,551 SNP sites covering loci selected by each clinical testing laboratory and the 96-SNP PCR panel. As a consequence, the genotypes identified utilizing the 96-SNP PCR panels are expected to be recapitulated in the NGS sequencing data. Custom oligonucleotide capture-enrichment panels were synthesized using Nimblegen (BCM-HGSC-CL) or Illumina (Broad CRSP). DNA sequencing was on the Illumina HiSeq 2500 platform. The performance of the two capture platforms was harmonized and applied to 14,515 (BCM-HGSC-CL) and 10,500 (LMM/Broad) samples. Both sites reported DNA sequence coverage of >99.9% of the targeted bases in these studies.

## Results

### Validation of the 96-SNP PCR Panel Performance

The BCM-HGSC-CL and LMM/Broad laboratories each utilized 96-SNP PCR panels to genotype and track DNA from the 25,015 samples collected at 11 clinical sites, throughout the process of clinical DNA sequencing and the development of clinical diagnostic DNA reports for participants in the NIH eMERGE III program. The two clinical testing sites utilized the same analytical platform foundation, employing slightly different SNP sites for the assays, but generally similar workflows (Figure 1), to test for concordance between data generated from the 96-SNP PCR panel genotyping and the ultimate DNA sequence data. The 96-SNP PCR panel platform was first shown to generate reliable genotyping data in validation experiments, testing 26 DNA control samples. In those validation experiments, the SNP call rates ranged between 96.8-100% depending on the sex of the sample. Inter and intra-run reproducibility was 100% for these 26 control DNA samples, in three independent rounds of tests.

**Figure 1.**
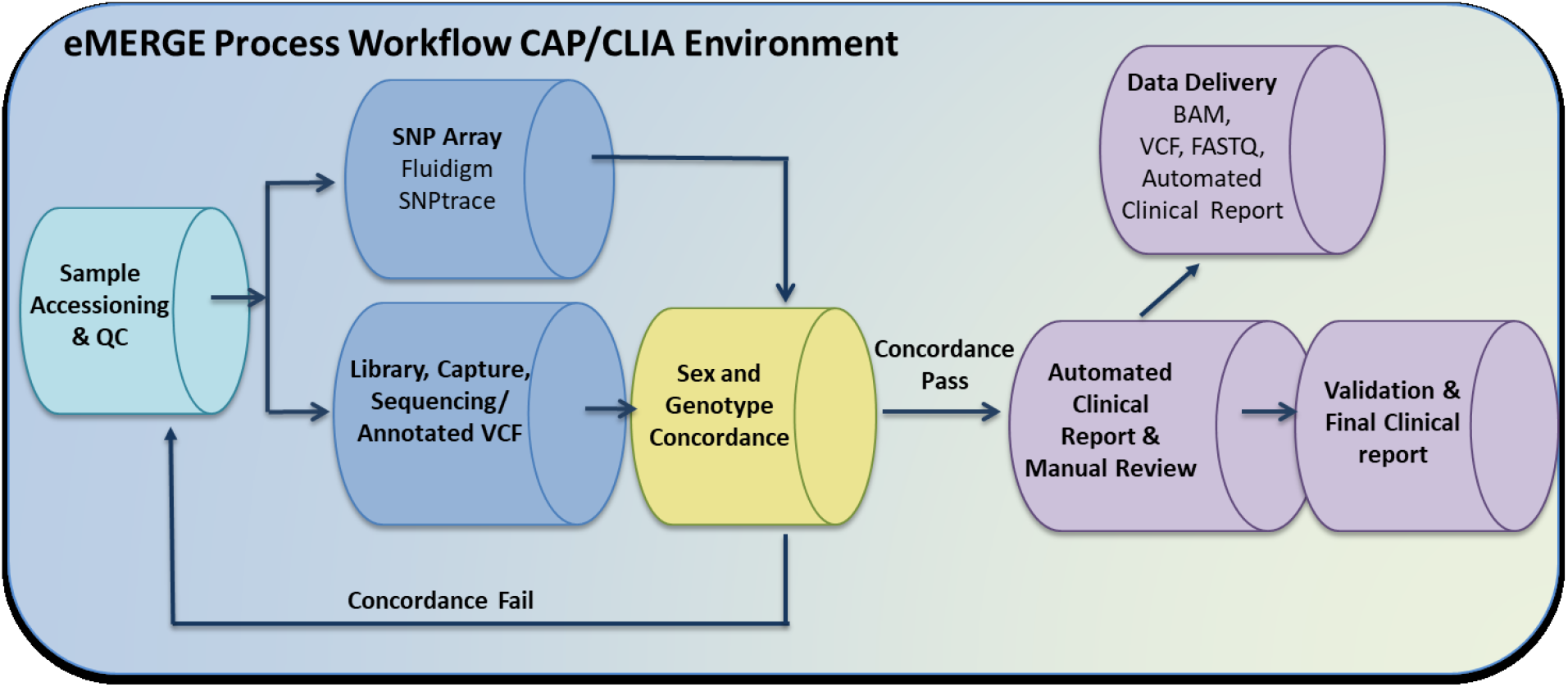
eMERGE sample processing workflow. Steps indicating where aliquots of DNA are taken from samples that are presented to the Clinical DNA Sequencing Laboratory for accession, to test via the Fluidigm 96-SNP PCR panel assay. Data from the Fluidigm 96-SNP PCR panel assay are compared with DNA sequence data from the DNA sequencing pipeline as a quality control step, ahead of the Automated Clinical Reporting step.

The 96-SNP PCR panel platform was applied for tracking the 25,015 eMERGE III test samples at the two clinical laboratory sites, during routine data production. The average SNP call rates were 97.3% and 97.5% for samples processed at the BCM-HGSC-CL and the LMM/Broad, respectively. A comparison of the 96-SNP PCR panel platform data obtained upon sample accession at each testing laboratory, with the subsequent sequence data, showed that there were no ‘sample swaps’ that had occurred during test laboratory processing.

### Identification and Technical Validation of Sex Discrepancies

The two testing laboratories utilized slightly different workflows to identify and then technically validate a total of 110 genetic vs. reported sex discrepancies from non-concordant data, when comparing the results from the 96-SNP PCR panel to the reported sex at the time of sample accession, for the 25,015 eMERGE III clinical samples (0.44%).

At the BCM-HGSC-CL, of the 14,515 samples processed, 73 samples with initial data suggesting discrepancies were retested with the same 96-SNP PCR panel to identify technical errors. Identical results were obtained for 70 of the re-tested samples (Table 1). For the remaining 3 cases, where the sex provided on test requisition was male, non-concordant or ambiguous data were obtained in both the initial and the repeated assays. For two of these samples, the automated software calls from one of each duplicate assay, indicated that the DNA source was from individuals with Klinefelter Syndrome (47, XXY). Review of the SNP scatter plots for these samples, however, including careful examination of the data from SNPs located on autosomes, indicated that these inconsistent sex calls most likely resulted from sample contamination involving a mixture of male and female DNAs (Figure 2).

**Table 1.**
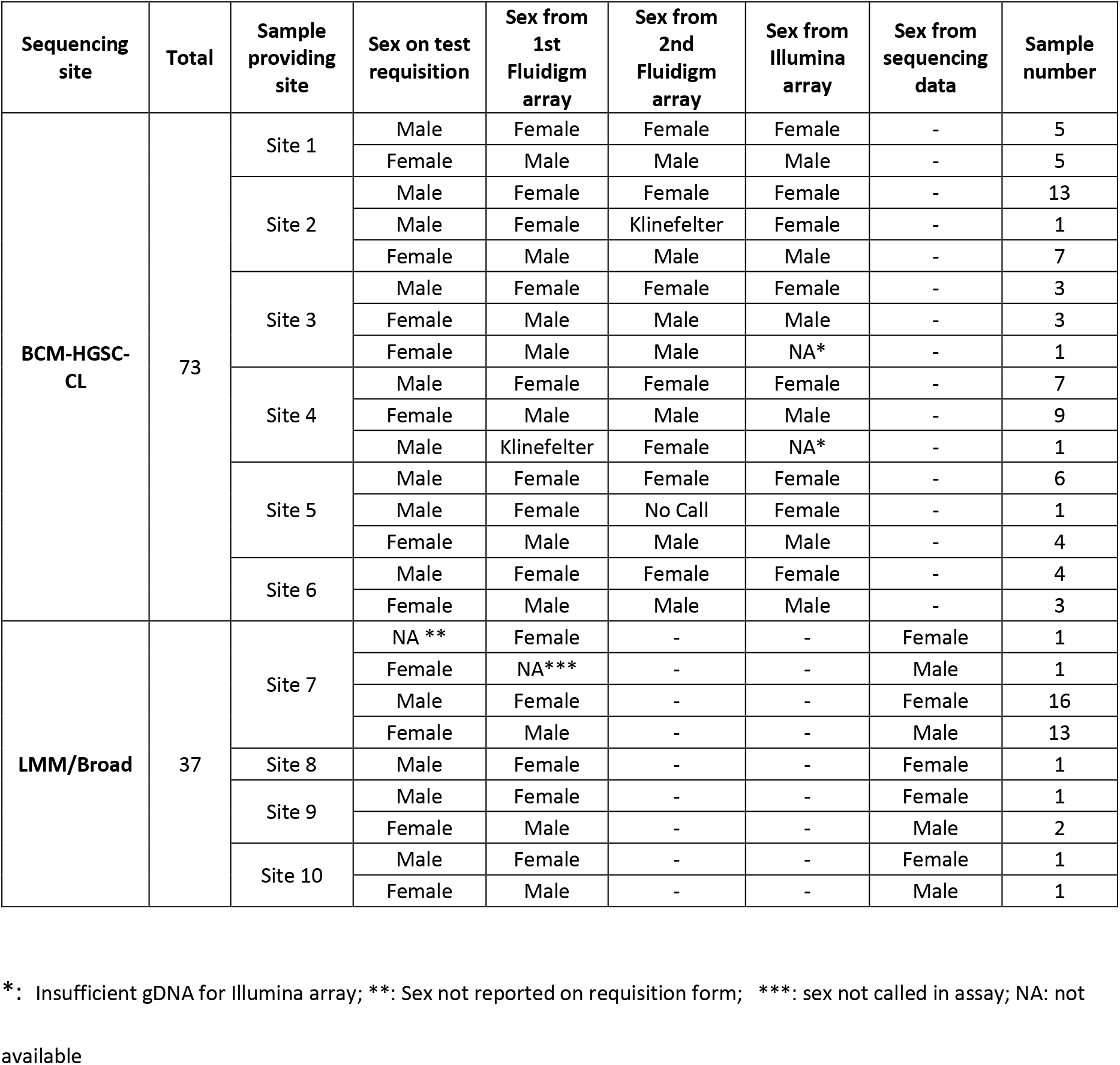
Comparison of genetic sex determined in various assays and reported sex on test requisition.

**Figure 2.**
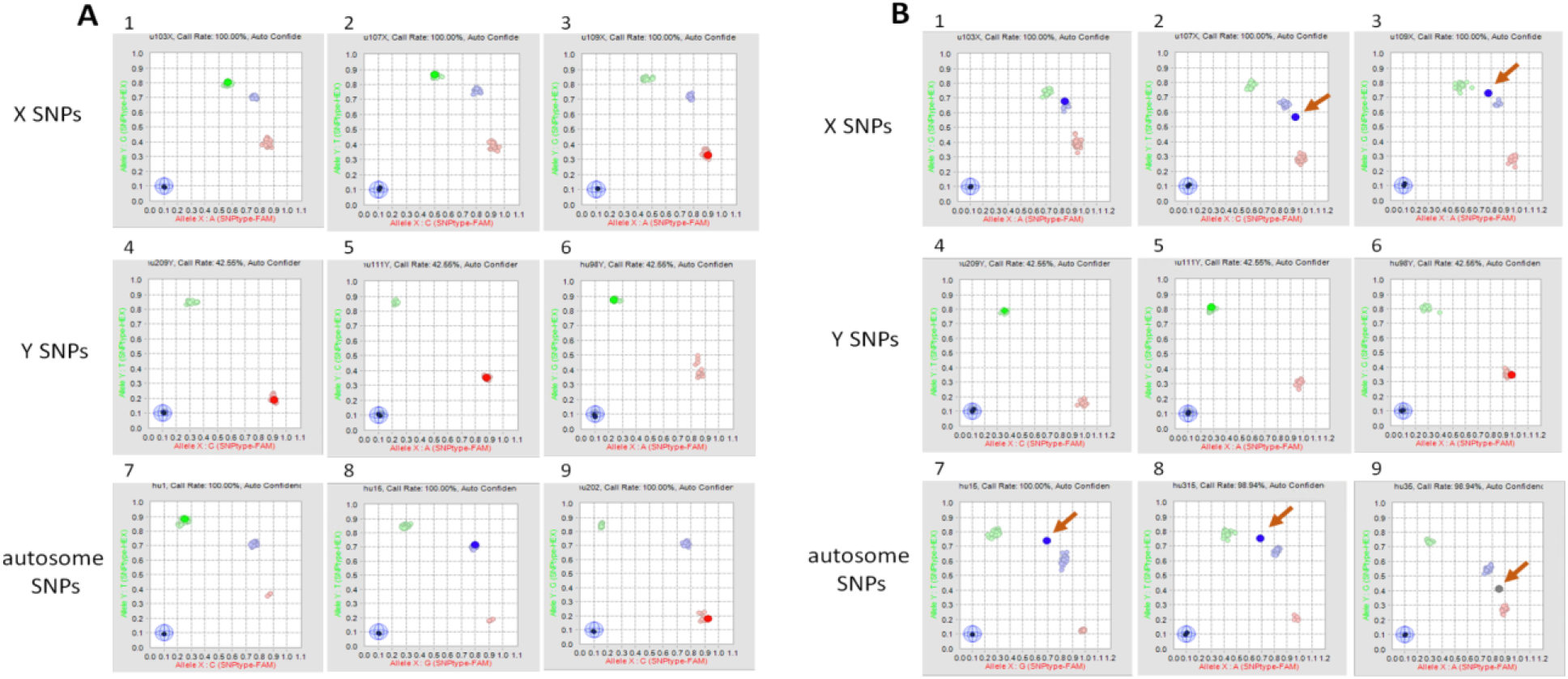
Scatter Plot Analysis of 96 SNP PCR Panel Reveals Sample Contamination. Scatter plot analysis from vendor software, showing a normal DNA male sample (A) or a contaminated sample containing a mixture of male and female DNAs (B). Panel 1-3: SNPs on X Chromosome; Panel 4-6: SNPs on Y Chromosome; Panel 7-9: autosomal SNPs. Each panel shows the data from a single SNP, as compared to clusters from all other SNPs. Clusters are shown as either homozygous (red or green), or heterozygous (blue) positions. In Panels B2, 3, 7-9, single SNPS are represented as outside the expected (arrows) resulting in erroneous or ‘no-call’ from the software.

Next, Illumina HumanCore Exome Arrays were utilized as an orthogonal high-density hybridization genotyping assay to further test 71 of the 73 samples with inconsistencies between the 96-SNP PCR panel genotyping assay and the initially reported sex at test requisition (two samples had insufficient genomic DNA available for the assay, including one of the suspected Klinfelter Syndrome cases). Genetic sex classifications from the data obtained using the hybridization method were identical to those from the 96-SNP PCR panel assay (Table 1). Among these, the data for one of the two suspected Klinefelter Syndrome samples, where adequate DNA was available, showed that the most likely explanation was indeed contamination of a female sample with additional male DNA.

At the Broad/LMM, site reported sex from the test requisition was compared with the genetic sex determined by both the 96-SNP PCR panel genotyping assay and the eMERGE III sequencing panel assay that was run in parallel with an independent DNA aliquot. Of the 10,500 samples processed at the Broad/LMM, 151 were initially identified as being discordant and were thus flagged for further investigation. The majority (95 samples) of the discrepancies were due to a technical failure of the Fluidigm 96-SNP PCR panel assay where no sex determination could be made. In all of these 95 samples, the sequencing sex matched the reported sex and no further action was taken. The remaining 56 of the 151 discordant samples had sex determination calls from both sequencing and genotyping assays. For 19 of the 56 samples, the sequencing and reported sex were concordant, but did not match the genotyping determined sex. Further review of these 19 samples showed that the genotyping assay calls were generally borderline or low confidence calls, suggesting sub-optimal performance of the 96-SNP PCR panel assay as the reason for the data discrepancy, rather than either a sex reporting error at accession or sample mix-up in the testing laboratory. The remaining 37 samples had highly confident sex determination calls from both the 96-SNP PCR panel assays and the subsequent DNA sequencing that were concordant, but did not match the site reported sex. These 37 samples were flagged for additional investigation with the clinical sites, in an effort to reconcile the difference between the reported and genotype determined sex.

### Origins of the sex discrepancy

The sample mix-ups that had resulted in the 110 discordant genotype-predicted sex with their reported sex at accession had not occurred within the clinical DNA sequencing laboratories. This was achieved by comparing the genotyping of the 96-SNP SNP panel to the sequencing data. These results indicate that most errors were introduced prior to shipment of samples to the sequencing centers. Clinical sampling sites were therefore contacted to further investigate the cause of the sex discrepant samples. Local handling errors were identified from test requisitions, sample extraction, and sample handling procedures for 54 cases. Forty-six of these had information that was incorrectly or incompletely entered on the test requisitions and were resolved by examination of other records. In 6 other cases it was determined that incorrect samples had been shipped from the sampling sites to the genome centers. Biological explanations for the discrepant tracking data were identified for an additional 12 cases. In four of these 12 cases, further examination of records revealed that the samples were provided by transgender participants. In addition, 8 sex discrepant samples were determined to be from individuals who had received stem cell or bone marrow transplants. Causes of the sample genetic vs. reported sex discrepancy are listed in Table 2.

**Table 2.**
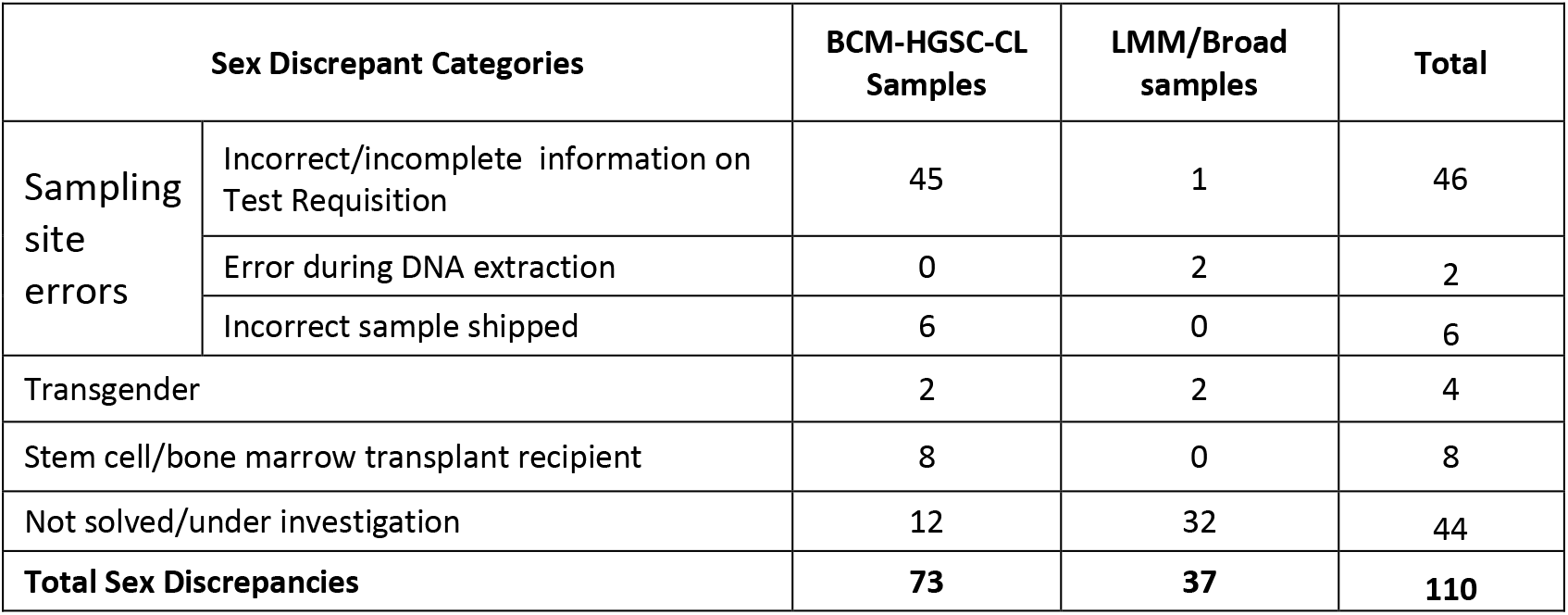
Causes of sample sex discrepancy.

Where possible, the information on test requisition forms were amended and correct clinical reports were issued for 44 cases processed at the BCM-HGSC-CL, or the incorrect samples were replaced and re-processed. Twelve cases sequenced at the BCM-HGSC-CL with sample-mix ups due to unknown causes were withdrawn from the study. Similarly, 32 unsolved cases sequenced at LMM/Broad were either withdrawn, or remain under investigation.

## Discussion

Sample tracking errors in clinical genetic diagnostic laboratories may have significant consequences for patients, leading to unnecessary surgical or diagnostic procedures, or delaying proper treatments. Additional consequences include inflated costs and diminished confidence in performance of the testing laboratories. To discover sample assignment errors during the processing of 25,015 clinical samples in the eMERGE III program, two clinical DNA sequencing laboratories utilized a Fluidigm-based 96-SNP PCR panel assay to genotype 96 polymorphic SNPs in aliquots of DNA in new shipments from participant enrollment sites. Data from the 96-SNP PCR panel assay were later compared to DNA sequences derived from the same samples, after they had been processed through the complex laboratory workflows. These analyses enabled a determination of whether the two data sets matched, indicating no sample swaps had occurred between sample arrival and DNA sequencing during the analytic phase. This overall robustness of sample tracking at the two clinical DNA sequencing laboratories reflects the mature automation and LIMS management that was applied to precisely track this volume of samples through the sequencing processes within the clinical testing laboratories.

In this study, we expanded the application of the Fluidigm platform based 96-SNP PCR panel genotyping assay to assess sample tracking fidelity during the pre-analytical (pre-testing) phase, before DNA samples arrived at the testing laboratories. The methodology relied upon a comparison of the sex inferred by the 96-SNP PCR panel genotyping assay to the recorded sex of each participant, who provided a sample for DNA extraction at test collection sites. The analyses identified 110 instances where the genotypes from the 96-SNP PCR panel assay did not represent the pattern expected from the sex of the participant that was assigned at the time of the test requisition. When the results were repeated using the same methodology, 70 samples yielded the exact same result. Three exceptions, that consistently provided ambiguous results, were samples where inherent sex chromosome imbalances were not expected to be resolved by the Fluidigm 96-SNP PCR panel assay. Other non-concordant samples were confirmed by combinations of tests with the initial Fluidigm 96-SNP PCR panel assay, array genotyping or by the eMERGE III DNA sequencing sites on the X-chromosome. Together these data provided high technical assurance that there were 110 instances of tracking anomalies within the sample set and that these likely arose before the materials were received at the clinical DNA sequencing laboratories suggesting quality control needs to focus on the preanalytical phase of the process.

The origin of the sample tracking errors at sample collection sites were determined in 66 of the 110 cases (60%), while leaving the remaining 44 cases unsolved and under investigation. Of these 66, the largest category was represented by the 54 cases that were the result of clerical or shipping errors (81%). Addressing these issues resulted in protocol alterations and establishment of procedures to promote more robust future handling. The remaining 12 cases (18% of the 66 solved) had biological underpinnings to the discordant results, as 8 were due to stem cell/bone marrow transplants while 4 were from transgender individuals. Future sample gathering procedures should be adjusted to ensure that participants are invited to note either of these prior events at an appropriate time, so that they are available for quality control.

The overall level of genetic and reported sex discordance of 0.44% is likely an underestimate of the true error rate in this study, as the misclassification of genetic sex from a random sample swap would be expected to result in incorrect, erroneous assignment, 50% of the time. The true ratio may be skewed by factors introducing a sex-bias in the direction of misclassification. This could be caused by skewed phenotypes of individuals with sex chromosome anomalies or that gender obfuscation may be socially driven in an unequal manner, depending on the gender identity of the individual. Overall, the rate is likely higher than the 0.44% identified here, but not anticipated to be higher than twice that level.

This level of tracking error is unacceptable for ongoing clinical practice, but the study does not represent the levels that will be expected in further clinical programs. Here we note that at least one laboratory declared their sample enrollments as ‘research samples’ and thus committed to later repeat assays under a fully compliant protocol, to verify any findings that may impact care. Others were able to quickly identify points of error and rectify their protocols to ensure faithful future sample handling. All sites committed to rechecking of records and reconciling actionable findings with orthogonal data, including family histories and biochemical tests, before returning results. The ‘lessons learned’ from these analyses ensure that a repeat of the same program would likely minimize any similar errors.

The samples that have a biological basis for sex discrepancies may still remain even if there were no clerical errors in a DNA testing program. In these cases, the data that are obtained through evaluation of genetic sex may be invaluable. For example, for an individual with a stem cell transplant, it would signal that blood cell changes may be reflective of the transplant-donor genotype and not necessarily relevant to the current host/participant. Similarly, individuals with sex chromosome anomalies may have been previously undiagnosed and the new information may prove beneficial.

The content of the 96-SNP PCR panel used here, was designed to provide genetic information about the identity, sex and quality of the human genomic DNA samples in this study(12). In the BCM-HGSC-CL, six of the 96 SNP sites in the 96-SNP PCR panel genotyping assay were linked to sex chromosomes, while at LMM/Broad 1 sex determining assay SNP covering AMG_3B gene is capable to differentiate male or female sample by 6 base pair insertion or deletion. The overall performance of the 96-SNP PCR panel at each site indicates that either choice of SNPs or indels can provide robust sex determination data. This panel has further utility and is capable of identifying closely-related samples and to pair tumor and matched normal samples. It may also detect mixed-sample contamination, although the precise levels of contamination may not be quantified. In our study, three samples showed that inconsistent sex calls in two rounds of 96-SNP PCR panel assay were due to sample contamination (Fig.2). For any samples with sex not called or called as Klinefelter Male, SNP array data were reviewed manually to determine if it was due to sample contamination.

In general, only 20 informative SNP loci are sufficient for unique individual sample identification(13, 14) and other SNP panels have been used for identification of human samples(6, 15, 16). A low-density QC genotyping array launched by Illumina which includes 15,949 markers has been utilized in genomic-based clinical diagnostics(17). Our studies showed that these two different SNP platforms exhibited consistent results when applied for sex identification. In comparison to the use of the Illumina Infinium array platform, the workflow for the 96-SNP PCR panel assay is faster (1 day workflow vs 3 day workflow) and more cost-effective. However, the Illumina Infinium array platform provides more information on linkage analysis, HLA haplotyping, ethnicity determination and other genetic information in addition to fingerprinting and thus may be preferred in some scenarios. Other commercial systems are also available to substitute for the platforms described here if they provide cost effective and precise data with similar qualities.

## Conclusions

In summary, the Fluidigm platform based 96-SNP PCR panel was successfully employed for sample tracking in a large network program with 11 sample collection sites and two clinical testing laboratories. The assay was robust when employed with slightly different SNP content and workflows and demonstrated zero sample-swaps within the clinical testing laboratories. When the assay data were explored to identify inconsistencies between genotype determined sex and the sex recorded at the time that participants enrolled for studies, 0.44% of samples were discordant. Errors were identified and rectified in procedures for sample recording, storage and shipping. In addition, transgender individuals and those with a history of transplant were identified, establishing a biological basis for the discrepancies. Although the number of sample tracking errors in the study is larger than can be detected by this assay, the findings established a lower limit for tracking accuracy and prompted revised methods of sample collection and accession, including computerized order entry and additional standardized automated sample processing.

## List of abbreviations

HGSC: Human Genome Sequencing Center
LMM: Laboratory for Molecular Medicine
NGS: Next generation DNA sequencing
STR: short tandem repeat
SNP: single nucleotide polymorphism
PCR: polymerase chain reaction
eMERGE: Electronic Medical Records and Genomics
EMR: electronic medical record
HGSC-CL: Human Genome Sequencing Center Clinical Laboratory
BCM: Baylor College of Medicine
CRSP: Clinical Research Sequencing Platform
STA: specific target amplification
LSP: locus specific primer
IFC: Integrated Fluidic Circuit
ASP: Allele Specific Primers
ACMG: American College of Medical Genetics
NHGRI: National Human Genome Research Institute
IRB: institutional review board

## Declarations

### Ethics approval and consent to participate

The Electronic Medical Records and Genomics (eMERGE) Network is a National Human Genome Research Institute (NHGRI)-funded consortium tasked with developing methods and best practices for utilization of electronic medical record (EMR) as a tool for genomic research. All 11 sample collection sites consented participants under institutional review board (IRB)-approved protocols and the two sequencing centers had IRB-approved protocols that deferred consent to the participating sites. The protocol number for Baylor College of Medicine was (#H-40455).

### Consent for publication

Not applicable.

### Availability of data and materials

Data are available in dbGaP for controlled public access (phs001616.v1.p1).

### Authors’ contributions

JH, HLR, RAG, DMM contributed to the study concept and design; JH, VK, HZ, MVH, CK, MES annotated and compiled information regarding sample accessioning; HZ, MVH, DM, EV performed NGS data analysis; NL, MES, GJ, HLR, RAG, DMM provided funding support for the project; Investigation: JH, VK, HZ, MVH, AM, SMH, CK, MES, AG, PS, MK, SB, LM, HLR, RAG, DMM conducted the research and investigation process of sample verification; AB, LZ, GC, DP performed the 96-SNP PCR panel and Illumina array genotyping assay; VK, CK, RR, KW, MM participated in the project administration; MES, AG, GJ, PS, MK, SB, CP provided eMERGE sample collections; JH, MM, EV, HLR, RAG, DMM supervised the studies; JH, HZ, MVH, HLR, RAG, DMM were the major contributors in original draft writing; JH, HZ, MVH, DM, AM, SMH, NL, RR, KW, AG, GJ, PS, MK, SB, MM, EV, EB, CP, LM, HLR, RAG, DMM participated in manuscript revision. All authors read and approved the final manuscript.

## Acknowledgments

We thank all eMERGE Phase III Network participants for their engagement in this research effort.

**EMERGE CONSORTIUM:**

Debra J Abrams^9^, Samuel E. Adunyah^14^, Ladia H. Albertson-Junkans^15^, Berta Almoguera^9^, Paul S. Appelbaum^16,17^, Samuel Aronson^3^, Sharon Aufox^7^, Lawrence J. Babb^5^, Adithya Balasubramanian^1^, Hana Bangash^18^, Melissa A. Basford^19^, Meckenzie Behr^9^, Barbara Benoit^20^, Elizabeth J Bhoj^9^, Sarah T. Bland^11^, Eric Boerwinkle^1,12^, Kenneth M. Borthwick^21^, Erwin P Bottinger^22,23^, Deborah J Bowen^24^, Mark Bowser^3^, Murray Brilliant^25^, Adam H. Buchanan^10^, Andrew Cagan^26^, Pedro J Caraballo^27^, David J Carey^28^, David S Carrell^15^, Victor M. Castro^26^, Gauthami Chandanavelli^1^, Rex L. Chisholm^7^, Rex L Chisholm^7^, Wendy Chung^29^, Christopher G Chute^30^, Brittany B. City^19^, Ellen Wright Clayton^19,31^, Beth L. Cobb^32^, John J Connolly^9^, Paul K Crane^33^, Katherine D. Crew^34^, David R. Crosslin^35^, Renata P da Silva^9^, Jyoti G. Dayal^6^, Mariza De Andrade^36^, Josh C. Denny^37^, Ozan Dikilitas^18^, Alanna J. DiVietro^19^, Kevin R. Dufendach^38,96^, Todd L. Edwards^19,39^, Christine Eng^2^, David Fasel^40^, Alex Fedotov^41^, Stephanie M. Fullerton^93^, Birgit Funke^42^, Stacey Gabriel^5^, Vivian S. Gainer^26^, Ali Gharavi^40^, Richard A. Gibbs^1^, Joe T Glessner^9,43^, Jessica M Goehringer^10^, Adam Gordon^7^, Adam S. Gordon^7^, Chet Graham^3^, Heather S Hain^9^, Hakon Hakonarson^9,43^, Maegan V. Harden^5^, John Harley^44,94^, Margaret Harr^9^, Steven M. Harrison^3,5^, Andrea L Hartzler^35^, Scott Hebbring^25^, Jacklyn N Hellwege^19,45^, Nora B. Henrikson^15,46^, Christin Hoell^7^, Ingrid Holm^47^, George Hripcsak^48^, Alexander L. Hsieh^48^, Jianhong Hu^1^, Elizabeth D. Hynes^3^, Gail P. Jarvik^8^, Darren K Johnson^10^, Laney K Jones^10^, Yoonjung Y Joo^49^, Sheethal Jose^6^, Navya Shilpa Josyula^50^, Anne E Justice^50^, Elizabeth W. Karlson^51^, Kenneth M. Kaufman^32,52^, Jacob M. Keaton^19,53^, Melissa A. Kelly^10^, Eimear E Kenny^54,55^, Dustin L Key^15^, Atlas Khan^56^, H. Lester Kirchner^50^, Krzysztof Kiryluk^40^, Terrie Kitchner^25^, Barbara J. Klanderman^3^, David C Kochan^18^, Viktoriya Korchina^1^, Christie Kovar^1^, Emily Kudalkar^3^, Benjamin R. Kuhn^57^, Iftikhar J. Kullo^18^, Iftikhar J Kullo^18^, Philip Lammers^14,58^, Eric B. Larson^15,59^, Matthew S. Lebo^3,60^, Ming Ta Michael Lee^10^, Niall Lennon^5^, Kathleen A Leppig^15,61^, Chiao-Feng Lin^3^, Jodell E. Linder^19^, Noralane M Lindor^62^, Todd Lingren^63,64^, Cong Liu^48^, Yuan Luo^65^, John Lynch^66^, Alyssa Macbeth^5^, Lisa Mahanta^3^, Bradley A. Malin^19^, Brandy M. Mapes^19^, Maddalena Marasa^56^, Keith Marsolo^67^, Elizabeth McNally^7^, Frank D Mentch^9^, Erin M. Miller^64,68^, Hila Milo Rasouly^56^, David Murdock^1,2^, Shawn Murphy^69^, Shawn N. Murphy^69^, Mullai Murugan^1^, Donna M. Muzny^1^, Melanie F. Myers^64,70^, Bahram Namjou^71^, Addie I Nesbitt^9^, Jordan Nestor^56^, Yizhao Ni^63,64^, Janet E Olson^62^, Aniwaa Owusu Obeng^72,73^, Jennifer A Pacheco^7^, Joel E Pacyna^74^, Divya Pasham^1^, Thomas N. Person^10^, Josh F. Peterson^19^, Lynn Petukhova^75,95^, Cassandra Pisieczko^10^, Siddharth Pratap^14^, Cynthia Prows^13^, Cynthia A Prows^13^, Megan J. Puckelwartz^7^, Alanna K Rahm^10^, James D Ralston^15,61^, Arvind Ramaprasan^15^, Luke V. Rasmussen^65^, Laura J. Rasmussen-Torvik^7,65^, Heidi L. Rehm^3,5^, Dan M. Roden^76^, Dan M. Roden^76^, Elisabeth A. Rosenthal^77^, Robb K. Rowley^6^, Maya S Safarova^18^, Avni Santani^9,78^, Juliann M. Savatt^10^, Daniel J Schaid^62^, Steven Scherer^1^, Baergen I. Schultz^6^, Aaron Scrol^15^, Soumitra Sengupta^48^, Gabriel Q Shaibi^79^, Ning Shang^48^, Himanshu Sharma^3^, Richard R. Sharp^74^, Richard R Sharp^74^, Yufeng Shen^48^, Rajbir Singh^14^, Patrick Sleiman^9^, Maureen E. Smith^7^, Jordan W. Smoller^80^, Duane T. Smoot^14^, Ian B. Stanaway^35^, Justin Starren^65^, Timoethia M. Stone^19^, Amy C Sturm^10^, Agnes S Sundaresan^81^, Peter Tarczy-Hornoch^35,82^, Casey Overby Taylor^10,83^, Lifeng Tian^9^, Sara L. Van Driest^84^, Matthew Varugheese^3^, Lyam Vazquez^9^, David L Veenstra^85,86^, Digna R. Velez Edwards^11,87^, Eric Venner^1^, Miguel Verbitsky^88^, Kimberly Walker^1^, Nephi Walton^10^, Theresa Walunas^49^,^89^, Firas H. Wehbe^65^, Wei-Qi Wei^11^,^19^, Scott T. Weiss^90^,^91^, Quinn S. Wells^92^, Chunhua Weng^48^, Chunhua Weng^48^, Ken L. Wiley, Jr.^6^, Marc S. Williams^10^, Janet Williams^10^, Leora Witkowski^3,42^, Laura Allison B. Woods^19^, Julia Wynn^29^, Lan Zhang^1^, Yanfei Zhang^10^, Hana Zouk^3,4^

^14^Meharry Medical College, Nashville, TN,

^15^Kaiser Permanente Washington Health Research Institute, Seattle, WA

^16^Department of Psychiatry, Columbia University, New York, NY

^17^NY State Psychiatric Institute, New York, NY

^18^Department of Cardiovascular Medicine, Mayo Clinic, Rochester, MN

^19^Vanderbilt University Medical Center, Nashville, TN

^20^Research IS and Computing, Laboratory for Molecular Medicine (LMM), Mass General Brigham, Cambridge, MA

^21^Hood Center for Health Research, Geisinger, Danville PA

^22^Hasso Plattner Institute for Digital Health, Icahn School of Medicine at Mount Sinai, New York, NY

^23^Division of Nephrology and Hypertension, Department of Medicine

^24^Department of Bioethics and Humanities, School of Medicine, University of Washington, Seattle, WA

^25^Marshfield Clinic Research Institute, Marshfield, WI

^26^Research IS and Computing, Laboratory for Molecular Medicine (LMM), Mass General Brigham, Cambridge, MA

^27^Department of Medicine, Mayo Clinic, Rochester, MN

^28^Molecular and Functional Genomics, Geisinger, Danville PA

^29^Department of Pediatrics, Columbia University Medical Center, New York, NY

^30^Schools of Medicine, Public Health, and Nursing, Johns Hopkins University, Baltimore, MD

^31^Center for Biomedical Ethics and Society, Vanderbilt University, Nashville, TN

^32^Cincinnati Children’s Hospital Medical Center, Cincinnati, OH

^33^Department of Medicine, School of Medicine, University of Washington, Seattle, WA

^34^Department of Medicine and Epidemiology, Columbia University, New York, NY

^35^Department of Biomedical Informatics and Medical Education, University of Washington, Seattle, WA

^36^Department of Health Science Research, Division of BioStatistics and Informatics, Mayo Clinic, Rochester, MN

^37^All of Us Research Program, National Institutes of Health, Bethesda MD

^38^Divions of Neonatology and Biomedical Informatics, Cincinnati Children’s Hospital Medical Center, Cincinnati, OH

^39^Division of Epidemiology, Department of Medicine, Vanderbilt University Medical Center, Nashville, TN

^40^Department of Medicine, Columbia University, New York, NY

^41^Irving Institute for Clinical and Translational Research, Columbia University, New York, NY

^42^Harvard Medical School, Boston, MA

^43^Department of Pediatrics, University of Pennsylvania School of Medicine, Philadelphia, PA

^44^Departments of Pediatrics and Medicine, University of Cincinnati College of Medicine, Cincinnati, Ohio

^45^Division of Genetic Medicine, Department of Medicine, Vanderbilt Genetics Institute

^46^Department of Health Services, School of Public Health, University of Washington

^47^Division of Genetics and Genomics and the Manton Center for Orphan Diseases Research, Boston Children’s Hospital, and the Department of Pediatrics, Harvard Medical School, Boston, MA

^48^Department of Biomedical Informatics, Columbia University, New York, NY

^49^Department of Medicine, Northwestern University Feinberg School of Medicine, Chicago, IL

^50^Population Health Sciences, Geisinger, Danville, PA

^51^Department of Medicine, Division of Rheumatology, Inflammation and Immunity, Brigham and Women’s Hospital, Boston, MA

^52^Cincinnati Veterans affairs

^53^Division of Epidemiology, Department of Medicine, Vanderbilt University Medical Center, Nashville, TN

^54^Institute for Genomic Health, Icahn School of Medicine at Mount Sinai, New York, NY

^55^Departments of Medicine and Genetics and Genomic Sciences, Icahn School of Medicine at Mount Sinai, New York, NY

^56^Division of Nephrology, Department of Medicine, Vagelos College of Physicians & Surgeons, Columbia University, New York, NY

^57^Pediatric Gastroenterology & Nutrition, Geisinger, Danville, PA

^58^Baptist Cancer Center, Memphis, TN

^59^Division of General Internal Medicine, University of Washington, Seattle, WA

^60^Brigham and Women’s Hospital, Harvard Medical School, Boston, MA

^61^University of Washington Biomedical and Health Informatics, Seattle, WA

^62^Department of Health Sciences Research, Mayo Clinic, Rochester, MN

^63^Division of Biomedical Informatics, Cincinnati Children’s Hospital Medical Center

^64^College of Medicine, University of Cincinnati, Cincinnati, Ohio

^65^Department of Preventive Medicine, Northwestern University Feinberg School of Medicine, Chicago, IL

^66^University of Cincinnati, Cincinnati, Ohio

^67^Department of Population Health Sciences, School of Medicine, Duke University, Durham, NC

^68^Division of Cardiology, Cincinnati Children’s Hospital Medical Center, Cincinnati, Ohio

^69^Department of Neurology, Massachusetts General Hospital, Boston, MA

^70^Division of Human Genetics, Cincinnati Children’s Hospital, Cincinnati, Ohio

^71^Center for Autoimmune Genomics and Etiology, Cincinnati Children’s Hospital Medical Center (CCHMC), Cincinnati, Ohio

^72^The Charles Bronfman Institute for Personalized Medicine, Icahn School of Medicine at Mount Sinai, New York, NY

^73^Departments of Pharmacy, Medicine and Genetics and Genomic Sciences, Icahn School of Medicine at Mount Sinai, New York, NY

^74^Biomedical Ethics Research Program, Mayo Clinic, Rochester, MN

^75^Department of Epidemiology, Mailman School of Public Health, Columbia University, New York, NY, USA

^76^Departments of Medicine, Pharmacology, and Biomedical Informatics, Vanderbilt University Medical Center, Nashville, TN

^77^Division of Medical Genetics, School of Medicine, University of Washington, Seattle, WA

^78^Department of Pathology and Laboratory Medicine, University of Pennsylvania Perelman School of Medicine, Philadelphia, PA

^79^Center for Health Promotion and Disease Prevention, Arizona State University, Phoenix, AZ

^80^Department of Psychiatry and Center for Genomic Medicine, Massachusetts General Hospital

^81^Population Health Sciences, Geisinger, Danville, PA

^82^Department of Pediatrics (Neonatology), University of Washington, Seattle, WA

^83^Department of Medicine, Johns Hopkins University, Baltimore, MD

^84^Departments of Pediatrics and Medicine, Vanderbilt University Medical Center, Nashville, TN

^85^Department of Pharmacy, University of Washington, Seattle, WA

^86^The Comparative Health Outcomes, Policy & Economics (CHOICE) Institute, Seattle, WA

^87^Department of Obstetrics and Gynecology, Division of Quantitative Sciences, Vanderbilt University Medical Center, Nashville, TN

^88^Division of Nephrology, Department of Medicine, Columbia University, New York, NY

^89^Center for Health Information Partnerships, Northwestern University, Chicago, IL

^90^Department of Medicine, Brigham and Women’s Hospital, Harvard Medical School, Boston, MA

^91^Department of Medicine, Harvard Medical School, Boston, MA

^92^Division of Cardiovascular Medicine, Department of Medicine, Vanderbilt University Medical Center, Nashville, TN

^93^Department of Bioethics and Humanities, School of Medicine, University of Washington, Seattle, WA

^94^Center for Autoimmune Genomics and Etiology (CAGE), Cincinnati Children’s Hospital Medical Center, Cincinnati, Ohio

^95^Department of Dermatology, Vagelos College of Physicians & Surgeons, Columbia University, New York, NY

^96^Department of Pediatrics, University of Cincinnati, Cincinnati, OH

